# Motor signals modulate cortical but not subcortical processing of self-initiated sounds

**DOI:** 10.64898/2026.07.10.737812

**Authors:** Laura Raiff, Gabrielle Butler, Kailyn McFarlane, Bharath Chandrasekaran, Kevin R. Sitek

## Abstract

When we produce sounds ourselves, the brain modulates the auditory neural response through an efference copy mechanism, allowing us to distinguish between self-initiated and externally generated auditory inputs. However, the precise level of the auditory pathway at which this attenuation occurs remains unclear. While evidence from animal models suggests that early auditory processing of self-generated sounds may be modulated by corticofugal signaling, localized cortical modulation would preserve the high-fidelity subcortical sound encoding while allowing flexible, context-dependent processing at higher levels. To probe potential motor influences in the early auditory system, we collected scalp-recorded frequency following responses (FFRs) from 33 normal-hearing adults during active (self-initiated) and passive (externally presented) listening conditions using a 170 ms speech stimulus. Data were collected with a vertical montage that emphasizes subcortical generators of the FFR. We observed no significant differences in the FFR between active and passive conditions in spectral power, response amplitude, pitch tracking, onset latency, or phase consistency. In contrast, cortical event-related potentials showed motor-induced suppression (MIS): reduced early peak amplitudes in the active condition after correcting for motor signals, increased phase consistency prior to auditory feedback, and more precise phase consistency at sound offset. In addition to indicating FFRs can be collected during a wider range of behavioral tasks without substantial motor contamination, our observation of the canonical MIS in cortical signals but not in FFRs suggests that MIS of self-initiated sounds primarily affects later stages of auditory processing rather than the early encoding reflected in the FFR.

## Introduction

Motor actions such as speaking or playing an instrument have auditory consequences. In this process, the motor system sends a copy of a motor command (efference copy) towards sensory areas, generating a prediction that encodes expected auditory consequences. This prediction then attenuates sensory neural responses to the expected self-generated auditory signals, likely freeing attentional resources from predictable self-generated sounds and preserving sensitivity to potentially meaningful external signals (Crapse & Sommer, 2008; Ford et al., 2014; von Holst & Mittelstaedt, 1950). Efference copy dysfunction is proposed to underlie auditory hallucinations in schizophrenia (Ford & Mathalon, 2005; Heinks-Maldonado et al., 2007), misophonia (Kumar et al., 2021), and inaccurate prediction coding during speech self-monitoring in neurodegenerative disease (Kim et al., 2023; Railo et al., 2020).

Predictive processing of auditory feedback is well-established in the cortex, where motor-induced suppression (MIS) is evident in auditory cortex responses to button-press-generated sounds, as measured in the N100 (Ford et al., 2014). Similarly, speech-induced suppression has been observed using magnetoencephalography (MEG; Houde et al., 2002; Niziolek et al., 2013), scalp-recorded electroencephalography (EEG; Ford & Mathalon, 2004; Sitek et al., 2013), and electrocorticography (ECoG; Chang et al., 2013; Flinker et al., 2010; Ozker et al., 2024). However, it remains unclear whether these modulating predictive signals propagate down the auditory pathway to subcortical structures like the inferior colliculus—which receives modulatory signals from auditory cortex (Suga et al., 2002; Winer et al., 1998)—and auditory brainstem nuclei, which reduce responses to self-generated sounds in animal models (Singla et al., 2017; Suga & Schlegel, 1972). The extent to which efference copy mechanisms influence earlier processing stages could significantly impact our understanding of the hierarchical organization of speech motor control. This gap persists largely because previous work in humans has lacked a method that can record subcortical auditory function non-invasively.

The scalp-recorded frequency-following response (FFR) addresses this gap directly, offering a non-invasive window into subcortical auditory processing. Specifically, it captures synchronized neural activity across the auditory pathway in response to periodic acoustic stimuli (Chandrasekaran & Kraus, 2010; Coffey et al., 2019). The FFR waveform encodes sound with high fidelity, as measured by fundamental frequency tracking, harmonic representation, and phase consistency (Krizman & Kraus, 2019). Traditionally, the FFR was thought to have exclusively subcortical generators (Chandrasekaran & Kraus, 2010; Greenberg et al., 1987; Smith et al., 1975), but modern evidence suggests there are multiple generators across the auditory pathway (Coffey et al., 2016; Gnanateja et al., 2021). Still, subcortical sources remain important and often dominant contributors to the FFR (Bidelman, 2018; White-Schwoch et al., 2019).

We thus recorded FFRs to test whether efference copy mechanisms modulate sound encoding below the cortex, contrasting *passive* listening and *active* sound generation via button pressing. Using this button pressing paradigm, prior research in humans has shown significant attenuation of cortical responses to self-initiated compared to externally generated sounds (Baess et al., 2011; Martikainen et al., 2005). In animal models, attenuation has been observed at the subcortical level (Suga & Schlegel, 1972). Based on this evidence, we hypothesized that, in addition to the expected cortical effects, FFR metrics would be attenuated for self-initiated compared to passively presented sounds. If, however, we did not find differences in the FFR between active and passive conditions, this would suggest that MIS may primarily affect later stages of auditory processing rather than the early encoding captured by scalp-recorded FFRs, providing important evidence about the hierarchical organization of self-initiated sound processing along the central auditory pathway.

## Methods

### Participants

We collected EEG from 33 adults (24 female, 9 male) with typical hearing abilities. All participants demonstrated native or native-like fluency in English, reported no history of seizures or neurological conditions, and exhibited normal auditory thresholds defined as hearing sensitivity better than 25 dB HL at frequencies of 0.5, 1, 2, and 4 kHz bilaterally.

Participants ranged in age from 18 to 40 years old (mean = 25 years). Acquisition errors for two participants led to event code discrepancies and exclusion from our cortical analyses (n=31), but all participants (n=33) were included in the FFR analysis since these events were determined differently (see Experimental design below).

All experimental procedures received approval from the Institutional Review Board of Northwestern University (STU00219782), and participants provided written informed consent prior to engagement in any research activities.

### Experiment design

EEG was collected in response to a 170 ms /da/ synthetic speech token with a 100 Hz fundamental frequency (f0; Fig. 1a). The /da/ syllable was used because it elicits robust FFRs with clear harmonic and formant structure (Krizman & Kraus, 2019). Auditory stimuli were delivered binaurally via ER3C insert earphones (Etymotic Research, IL, USA), in alternating polarity, at approximately 75 dBA SPL. Stimulus presentation was administered through Presentation software (Version 23.0, Neurobehavioral Systems, Inc., Berkeley, CA, www.neurobs.com), with audio signals routed through a Fireface UC external sound card (RME Audio, Germany) to ensure precise temporal control and optimal audio quality.

**Figure 1.**
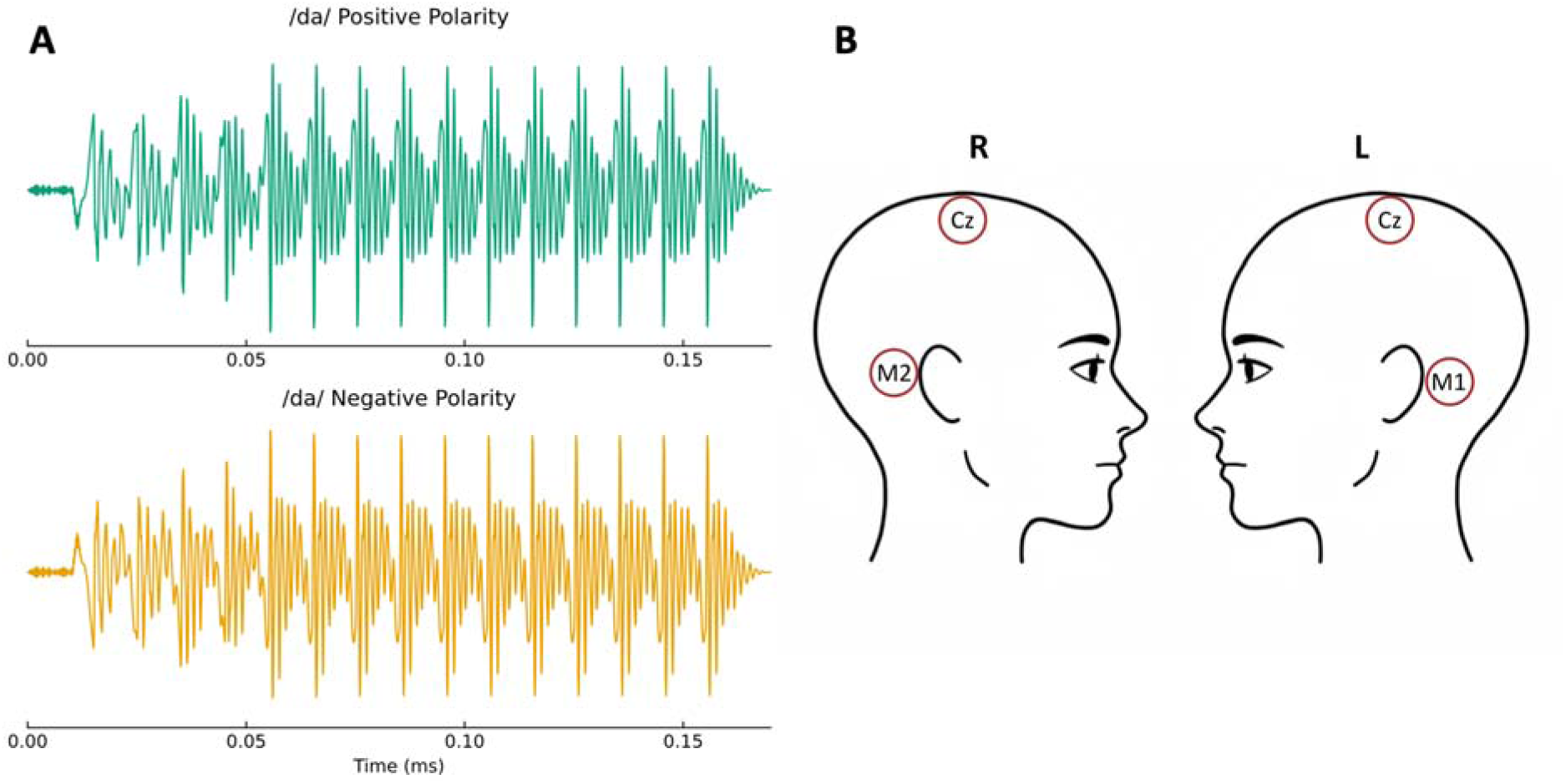
a) Waveform of audio stimulus /da/. Top waveform in green shows the positive polarity. Bottom waveform in orange shows the negative polarity. B) Vertical EEG montage used. M1 and M2 were used as reference channels for Cz.

The experiment comprised three distinct experimental conditions designed to differentiate between neural responses to self- versus externally generated auditory stimuli:

1. *Active*, in which participants initiated stimulus presentation by pressing a button at their own pace, targeting approximately 500 ms intervals. The target pace was demonstrated to them by the experimenter prior to the experiment;
2. *Passive*, in which the /da/ token was presented automatically (i.e., without button press initiation) in 500 ms intervals (on average; interstimulus intervals were jittered between 400–600 ms to mitigate anticipatory neural responses or entrainment to stimulus presentation rate); and
3. *Motor*, in which participants were instructed to press the button at the same pace as the other two conditions, but no auditory stimuli followed. This condition served as an experimental control to isolate neural correlates of the motor action of pressing the button.

Approximately 1200 trials were collected in each run of the active, passive, and motor conditions, allowing for 600 trials in each polarity. Individual experimental runs extended for approximately 10–15 minutes, with precise duration contingent upon the participant’s pace of button pressing in the active and motor conditions. Each participant completed 2400–6000 trials per condition, depending on available time, with 23 participants having at least 4800 trials per condition. Each session started with active, passive, and then motor trials, repeating that cycle at least twice. Participants were allowed to watch a silent, captioned show of their choosing to maintain a consistent level of arousal throughout testing (Coffey et al., 2019; Skoe & Kraus, 2010).

EEG data were acquired at a sampling rate of 16384 Hz using a BioSemi ActiveTwo system (Biosemi, Netherlands) via the ActiView software (version 9.02, www.BioSemi.com). A standard vertical electrode montage (vertex [Cz] linked to mastoids [M1 and M2]) was employed to optimally capture responses generated by more central brainstem structures (Fig. 1b; Galbraith, 1994). Motor responses were recorded via a physical-to-digital trigger interface (Triggy; Cortech Solutions, NC, USA), which transmitted synchronization markers to the Presentation software used to trigger audio playback during the active condition and to mark event onsets for the ERP analysis. For the active and passive conditions, the audio stimulus was simultaneously recorded through an analog input channel of the BioSemi ActiveTwo system. This channel was then used to identify precise event onset markers directly from the acoustic signal to define epochs for the FFR analysis. Data were acquired with open filters, so FFRs and cortical ERPs were measured from the same signal, depending on post-hoc filtering.

### FFR preprocessing analysis

FFRs were isolated by bandpass filtering the EEG data between 65–2000 Hz using a zero-phase FIR filter (MNE-Python default parameters). These data were then epoched starting 40 ms before the stimulus onset until 400 ms after, with -40 ms to 0 ms considered the baseline. To ensure data quality, an artifact rejection protocol was implemented to exclude epochs with a magnitude greater than 75 µV to remove contamination from eye blinks, muscular activity, or head movements. The filtered epochs were then averaged separately for each experimental condition (active and passive), resulting in condition-specific FFR waveforms for each participant. Due to the absence of audio stimuli, the FFR was not analyzed for the motor-only condition. All analyses used the polarity-added waveform.

To compare FFRs from active and passive conditions, we first visually compared the power spectra, computed using the Welch method over the vowel portion of the stimulus (50-150ms), and spectrograms to characterize neural response in each condition. For the time domain analysis, we computed root mean square (RMS) signal-to-noise ratios (SNR) for each participant’s responses, calculated by dividing the RMS of the response period by that of the baseline. RMS SNR quantifies the strength of the neural response compared to background neural activity. A higher SNR may reflect higher fidelity auditory processing (Bidelman, 2015; Krizman & Kraus, 2019). Additionally, we compared the latency of the onset of the FFR between conditions. The timing was determined by the primary and senior author who were blinded to group membership and the order trials was randomized. Intra-class correlation values were computed to assess reliability between raters (Liljequist et al., 2019).

For the frequency domain analysis, we extracted mean spectral power in the fundamental frequency range of the stimulus (90–110 Hz). Power values were averaged over the included time window (50–150 ms) and frequency band, resulting in one power value per condition per participant. We also computed phase consistency, derived from a short-time Fourier transform using a 40ms Hann window with 36 ms overlap, to quantify the degree to which the EEG signal time-locked across trials (Krizman & Kraus, 2019). Finally, we estimated *f*0 of the FFR for each participant and condition using an autocorrelation function and finding the lag with the highest correlation value.

### Cortical ERP processing and analysis

Differences between self-initiated and passively presented sounds have been reported previously with cortical ERPs (Ford et al., 2014), but not with FFRs. To validate that our data collection scheme was sensitive to any self-initiated vs. passively presented auditory processing differences, we computed cortical ERPs from the same data using a zero-phase FIR bandpass filter (1–30 Hz; MNE-Python default parameters). Remaining preprocessing steps followed those used for FFR preprocessing.

To compare between conditions cortically, we computed the phase consistency and spectrograms as done with the FFR analysis.

To disentangle the motor and auditory responses, we subtracted the motor-only ERP from the active condition ERP within subjects. We compared the peak amplitudes between waveforms from the resulting subtraction to the passive condition ERP to assess for MIS of the auditory response.

All statistical analyses were conducted with a significance threshold of α = 0.05. All condition comparisons used paired t-tests. When multiple comparisons were assessed (phase consistency and f0 estimation), we used false discovery rate (FDR)-corrected q = 0.05. Post-hoc two one-sided tests (TOST) and Bayes Factor (BF) were computed for metrics that failed to reach significance. Computational analyses were executed in

Python, using MNE-Python framework. Processing and analysis code can be found on our GitHub repository: https://github.com/SoundBrainLab/EAM1-FFR. Our data is available online: doi:10.18112/openneuro.ds008161.v1.0.3

## Results

### FFR Results

We analyzed our FFR data using common metrics to assess differences between conditions. In Fig. 2a, we show the grand average, averaged across all participants, FFR for each condition, yielding two evoked responses (active and passive) per participant.

**Figure 2.**
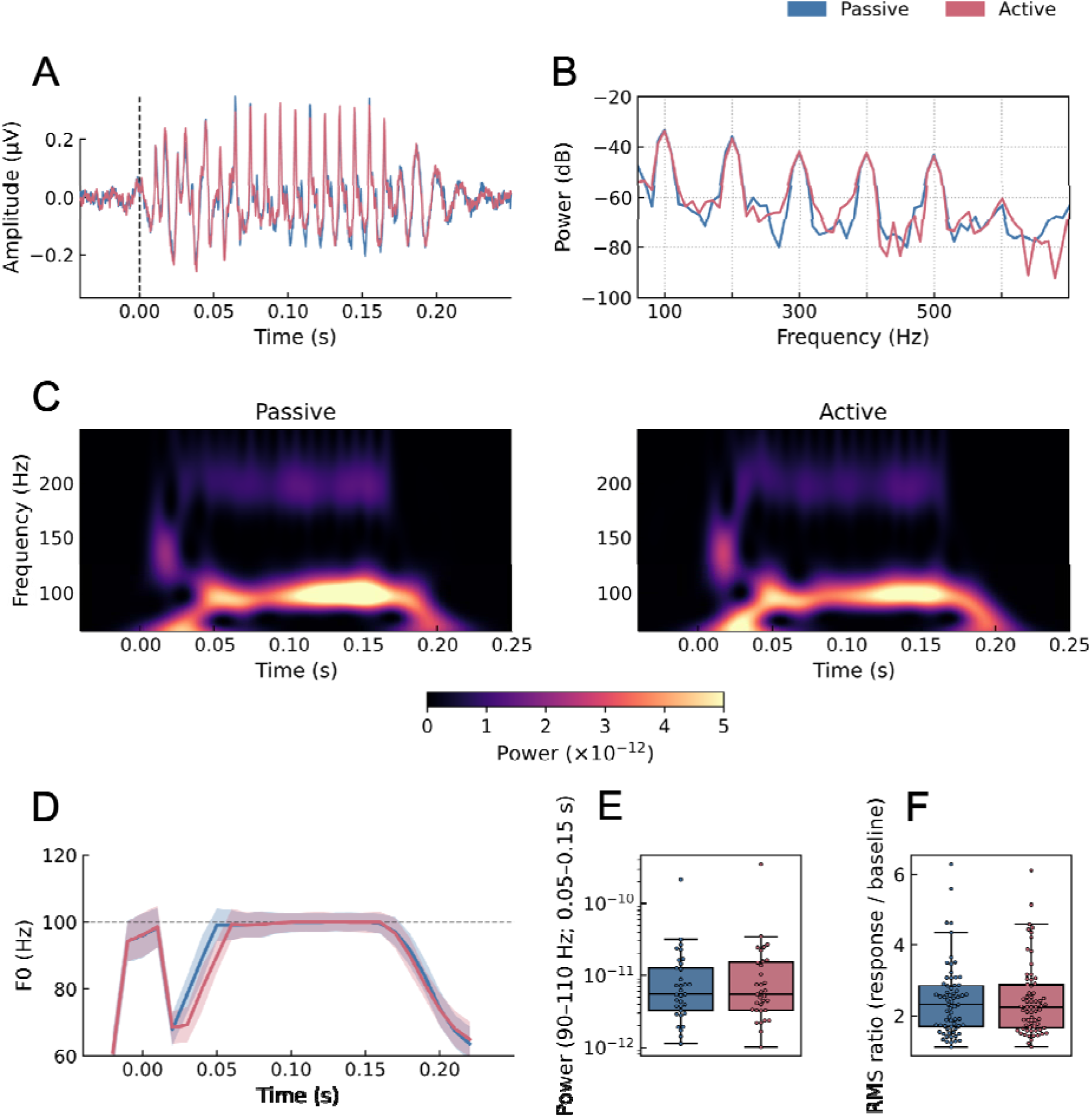
FFR Results for passive (pink) and active (blue) conditions. A: Grand average waveform. B: Power sp ctra. C: Spectrogram of FFRs from passive (left) and active (right) conditions. D: Pitch tracking over the time of the FFR. Stimulus pitch is indicated via gray dashed line at 100 Hz. E: Box plot comparing power around the f0 of the stimulus between passive and active conditions. F: Box plot comparing RMS ratio of the response:baseline between conditions.

In the power spectra, we observed clear peaks at the fundamental frequency (100 Hz) and its harmonics (200 and 300 Hz; Fig. 2b). Sustained energy at the fundamental frequency is shown throughout the vowel portion of the stimulus (50–150 ms) in both conditions’ spectrograms (Fig. 2c). Overall, the FFRs robustly encoded key frequency content of the stimulus.

Across metrics – including f0 tracking (Fig. 2d; q > 0.05), spectral power (Fig. 2e), RMS SNR (Fig. 2f), phase consistency (Fig. 3a; q > 0.05), and onset latency – no significant differences were observed between conditions (Table 1). For onset latency, the agreement between raters was moderate (ICC(A,1) = 0.664), and both agreed on excluding one subject. Overall, active trial responses started slightly, but not significantly, earlier. TOSTs were significant for both power and SNR metrics, indicating equivalency between passive and active conditions. Additionally, BF were highly favorable for supporting our null hypothesis given our data, although post-hoc tests for onset latency were inconclusive.

**Figure 3.**
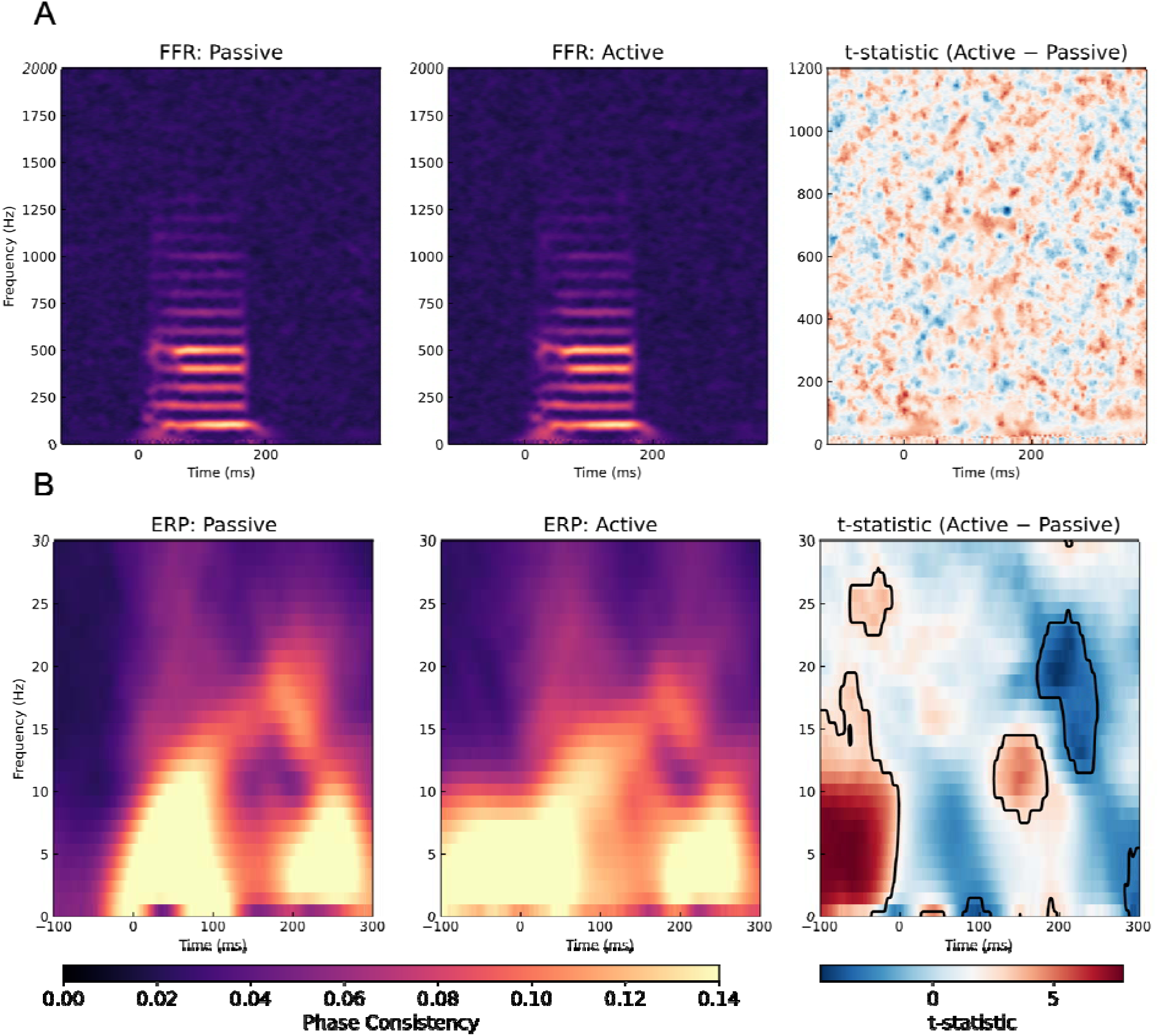
Phase consistency results. A: Phase consistency for passive (left) and active (middle) of the frequency-following responses (FFRs). There was no statistically significant difference between the conditions when comparing every pixel of the time-frequency map (right). B: Phase consistency for passive (left) and active (middle) of the event-related potentials (ERPs). There were several regions of statistical significance as enclosed by black lines on the t-statistic map (right).

**Table 1:** Statistical summary from frequency-following response (FFR) and cortical event-related potential (ERP) metrics. TOST = two one-sided tests (equivalency test); BF10 = Bayes Factor for alternative hypothesis; BF01 = Bayes Factor for null hypothesis.

Short title: Frequency-following responses to self-initiated sounds
| Metric | Passive Mean | Active Mean | Active-Motor Mean | t-value | p-value | TOST p-value | BF10 (BF01) |
| --- | --- | --- | --- | --- | --- | --- | --- |
| FFR Spectral Power | $9.70 \times 10^{-12}$ | $1.39 \times 10^{-11}$ | -- | -1.238 | 0.225 | < 0.05 | 0.34 (2.99) |
| FFR RMS SNR | 2.44 | 2.45 | -- | -0.167 | 0.868 | < 0.001 | 0.14 (7.30) |
| FFR Onset Latency (ms, rater averaged) | 13.7 | 12.5 | -- | -1.547 | 0.132 | 0.113 | 0.60 (1.79) |
| Cortical ERP Peak Amplitude (μV) | 3.92 | -- | 3.07 | 2.436 | < 0.05 | -- | 2.39 (0.42) |

### Cortical ERP results

While FFRs did not appear to differ between active and passive conditions, previous studies found cortical ERP differences (Baess et al., 2011; Ford et al., 2014; Martikainen et al., 2005). We therefore calculated cortical ERPs using a 1–30 Hz bandpass filter and compared these responses to actively and passively generated sounds (Fig. 4a).

**Figure 4:**
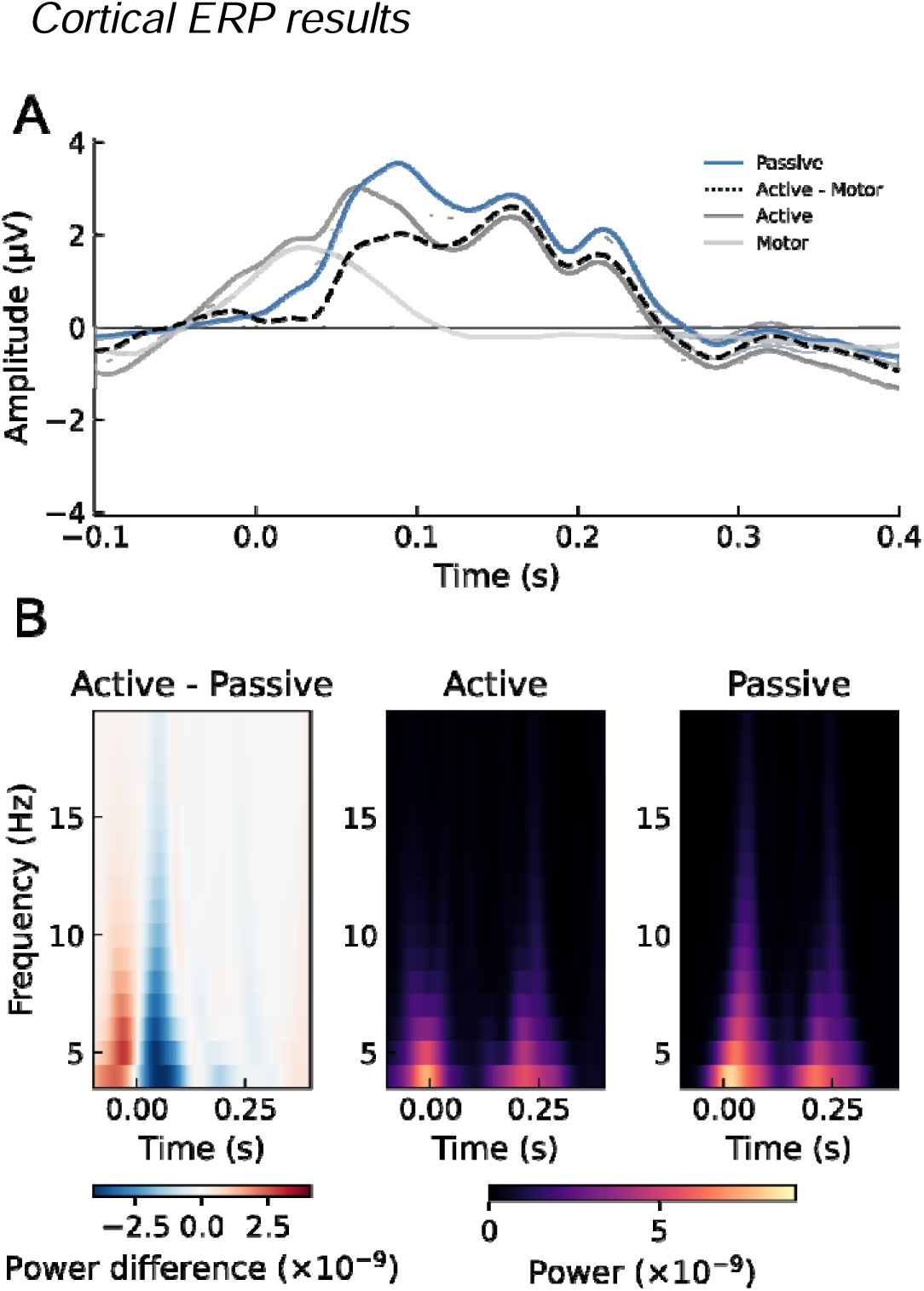
ERP results. A: Waveform of the active (dark gray), passive (blue), motor (light gray), and motor-corrected active (dashed line) conditions. Event onset is shown via the dashed gray line. 95% confidence intervals are shown as shading. B: Spectrograms showing the power of frequencies between 1 and 30 Hz across the time of the waveforms for difference between conditions (active - passive; left), active (middle) and passive (right) conditions.

The active condition had statistically significantly greater phase consistency relative to the passive condition particularly around the offset of the sound (approx. 170ms; Fig. 3b). Passive trials also had a period of significantly increased phase locking post-stimulus, but this appears to occur later than active trials. (See Table 2 for a description of regions of increased phase consistency.) In both active and passive, there was increased power around sound onset and offset (Fig. 4b). Although there was no significant difference, the active condition demonstrates higher earlier power before sound onset likely due to the motor contributions (Fig. 4b, left).

**Table 2:** Summary of statistically significant clusters of increased phase consistency in cortical responses (Figure 3B).

| Trial Type | Peak Frequency (Hz) | Peak Time (ms) | Peak t-value | Frequency range (Hz) | Time range (ms) |
| --- | --- | --- | --- | --- | --- |
| Active | 5 | -62.6 | 7.910 | 0 – 19 | -100 – -1 |
| Active | 11 | 150.4 | 5.327 | 8 – 14 | 117 – 189 |
| Passive | 20 | 199.2 | -4.609 | 12 – 24 | 170 – 248 |

Finally, to disentangle the auditory and motor contributions to the active condition ERP, we subtracted the ERPs from the motor-only ERP from the active ERP for each participant. The resulting subtracted waveform resembled the passive (pure auditory) ERP (Fig. 4a). The motor-corrected active ERP had a statistically significant reduced amplitude compared to the passive condition, indicative of MIS of the auditory processing. See Table 1 for a statistical summary.

## Discussion

In this study, we investigated the neural processing differences between self-initiated and externally presented sounds by comparing scalp-recorded electrophysiological responses during active and passive listening conditions. Using EEG to record neural responses to a /da/ syllable in 33 participants, we found no statistically significant differences in FFR metrics (spectral power, RMS SNR, *f*0 tracking, onset latency, or phase consistency) between conditions. In contrast, cortical ERPs showed clear modulation: active trials exhibited greater phase consistency before trial onset and during trial offset, and motor-corrected active trials demonstrated suppressed peak amplitudes compared to passive trials. This dissociation between subcortical and cortical responses provides evidence that efference copy signals may primarily influence later stages of the auditory pathway.

The motor-corrected ERP showed a significant reduction in peak amplitude relative to the passive condition. Subtracting the motor component rules out motor-evoked activity overlapping with the auditory ERP, instead indicating motor-induced auditory suppression, consistent with prior button-pressing work (Baess et al., 2011; Ford et al., 2014; Jack et al., 2021; Knolle et al., 2013).

We also observed efference copy mechanisms acting on the phase of cortical signals, described here as phase consistency but otherwise known as inter-trial phase coherence, phase-locking factor, or neural synchrony. Active ERPs had higher phase consistency prior to sound onset, while passive ERPs displayed minimal phase consistency pre-stimulus. Because cortical signals were epoched to acoustic onsets, which followed the button press by only tens of milliseconds, the pre-stimulus window captures activity occurring during preparation and shortly after motor execution but before auditory feedback. Increased neural synchrony observed in concordance with MIS has been previously reported in vocalization (Chen et al., 2011; Ford et al., 2007), visually cued button pressing (Popovych et al., 2016), and in rodents prior to exploratory whisking (Hamada et al., 1999). In line with the previous interpretations, the increased pre-stimulus synchrony likely acts as a predictive mechanism given its association with subsequent MIS, resetting the phase of populations of neurons in anticipation of a self-initiated sensory stimulus. We also observed increased post-stimulus phase consistency in both conditions. However, active trials appeared to track the offset of the stimulus with higher precision, as passive trial consistency was seen at a noticeable delay from the true offset. Thus, this prediction mechanism may persist beyond stimulus onset to track the stimulus offset with temporal precision.

Evidence for enhanced subcortical auditory phase consistency has only been observed in externally driven paradigms, such as rhythmic entrainment or stimulus-driven synchronization, where auditory input continuously constrains motor timing (Tierney & Kraus, 2013). Our study instead involved volitional, self-initiated actions. The lack of FFR modulation here may therefore suggest that motor-related timing signals alone are insufficient to alter subcortical phase-locking, and that externally imposed temporal constraints are necessary. One possible explanation is that different forms of motor engagement (entrainment vs self-initiation) recruit distinct neural systems that differ in their interactions with the auditory pathway (Krieghoff et al., 2011; Thaut et al., 2015). Future work should more directly examine how the source and nature of motor signals influences where along the auditory pathway motor integration occurs.

The null FFR results also raise a broader question about the extent to which cognitive processes (e.g., volitional movement or attention) can influence subcortical auditory processing. Descending corticofugal projections anatomically support top-down modulation of early auditory encoding (Winer, 2005; Winer et al., 1998), but to our knowledge no study has examined whether volitional motor control represents one such influence on the FFR. Selective attention, a related cognitive process, has received considerably more investigation in this context, though with conflicting results. Some studies have reported enhanced FFR amplitude during directed auditory attention (Galbraith et al., 2003; Price & Bidelman, 2021) or suppression during audiovisual presentation of speech (Musacchia et al., 2006), while others have found no systemic FFR modulation (Stoll et al., 2025; Varghese et al., 2015). Stoll et al. (2025) used naturalistic speech and measured responses spanning the full auditory pathway (from the auditory nerve through the brainstem and cortex), and concluded, consistent with our volitional movement results, that selective attention only influences neural encoding in the cortex. Price and Bidelman (2021) found attentional effects on the FFR only at the source level, not at the scalp electrode level, suggesting that the sensitivity of the recording configuration may determine detectability of effects. Xie et al. (2018) found that attentional effects on the FFR were context dependent: decreased predictability reduced the fidelity of the stimulus encoding, though notably using a decoding approach potentially more sensitive to subtle modulation than conventional metrics used here. Importantly, our paradigm did not explicitly manipulate attention: participants were instructed to engage in passive listening across both active and passive conditions, making attentional differences between conditions an unlikely explanation for our results. Nevertheless, the methodological heterogeneity and limitations within the attention literature is informative for interpreting our volitional movement findings, as it highlights how recording approach and task design can shape whether top-down modulation is detected.

Overall, the lack of significant attenuation in the FFR during self-initiated sounds suggests that MIS may be more prominent at later stages of auditory processing, or undetectable in scalp-recorded FFRs under these experimental conditions. As Coffey et al. (2016; 2019) demonstrated, our electrode configuration (Cz referenced to linked mastoids) emphasizes peripheral and subcortical sources in the recorded FFR signal. Indeed, Galbraith et al. (2003)’s study found largest changes in FFR amplitude due to attention demands at electrodes farthest away from Cz, indicating that this topography reveals different generators and thus influences on the FFR. Neuroanatomically, direct projections from cortical motor areas to the auditory cortex could mediate suppression effects, while subcortical auditory structures may be less directly influenced by cortical motor signals that drive voluntary finger movement (Schneider et al., 2014). More broadly, predictive and cognitive influences on auditory processing may be selectively implemented rather than uniformly distributed across the auditory hierarchy. A more localized cortical modulation would preserve the high-fidelity encoding of sound at subcortical stages while allowing flexible, context-dependent processing at higher levels. Additionally, our null results should also reassure future FFR studies involving motor-related behavioral tasks that substantial motor contamination is unlikely.

An important limitation of the present study is that the motor task involved a simple button press rather than a naturally sound-generating behavior (e.g., speech). The efference copy signals engaged in this task may therefore differ from those involved in vocalization, limiting generalization to speech processing. Additionally, our short stimulus onset asynchrony of 500 ms precluded an N100 in the cortical results, limiting comparisons to previous literature reporting on cortical MIS. Finally, given evidence of sex differences in the FFR (Krizman et al., 2012), and our predominantly female sample (24 female vs. 9 male subjects), we were underpowered to assess sex effects and cannot rule out a female-biased response.

While human and animal studies demonstrated that motor signals directly affect auditory processing of self-initiated sounds, our findings suggest that the neural attenuation of self-initiated sounds is realized later in cortical stages rather than in early subcortical encoding primarily reflected in the FFR. Future studies using high-density EEG, source localization, or multimodal imaging approaches could better differentiate cortical and subcortical contributions, with cross-frequency coupling or connectivity analyses offering insights into how motor signals modulate the functional integration within the auditory pathway. Understanding how the brain differentiates self-initiated from external sounds has important implications for speech self-monitoring and auditory perception, and examining these mechanisms across development and in clinical populations (e.g., schizophrenia or Parkinson’s Disease) will further elucidate how the auditory system integrates motor and sensory information.

## Acknowledgments

We thank Amp Kangsumrith for assistance in collecting data and Gavin Bidelman for feedback on an earlier draft of this manuscript.

## References

Baess, P., Horváth, J., Jacobsen, T., & Schröger, E. (2011). Selective suppression of self-initiated sounds in an auditory stream: An ERP study. Psychophysiology, 48(9), 1276–1283. 10.1111/j.1469-8986.2011.01196.x

Bidelman, G. M. (2015). Multichannel recordings of the human brainstem frequency-following response: Scalp topography, source generators, and distinctions from the transient ABR. Hearing Research, 323, 68–80. 10.1016/j.heares.2015.01.011

Bidelman, G. M. (2018). Subcortical sources dominate the neuroelectric auditory frequency-following response to speech. NeuroImage, 175, 56–69. 10.1016/j.neuroimage.2018.03.060

Chandrasekaran, B., & Kraus, N. (2010). The scalp-recorded brainstem response to speech: Neural origins and plasticity. Psychophysiology, 47(2), 236–246. 10.1111/j.1469-8986.2009.00928.x

Chang, E. F., Niziolek, C. A., Knight, R. T., Nagarajan, S. S., & Houde, J. F. (2013). Human cortical sensorimotor network underlying feedback control of vocal pitch. Proceedings of the National Academy of Sciences, 110(7), 2653–2658. 10.1073/pnas.1216827110

Chen, C.-M. A., Mathalon, D. H., Roach, B. J., Cavus, I., Spencer, D. D., & Ford, J. M. (2011). The Corollary Discharge in Humans Is Related to Synchronous Neural Oscillations. Journal of Cognitive Neuroscience, 23(10), 2892–2904. 10.1162/jocn.2010.21589

Coffey, E. B. J., Colagrosso, E. M. G., Lehmann, A., Schönwiesner, M., & Zatorre, R. J. (2016). Individual Differences in the Frequency-Following Response: Relation to Pitch Perception. PLOS ONE, 11(3), e0152374. 10.1371/journal.pone.0152374

Coffey, E. B. J., Nicol, T., White-Schwoch, T., Chandrasekaran, B., Krizman, J., Skoe, E., Zatorre, R. J., & Kraus, N. (2019). Evolving perspectives on the sources of the frequency-following response. Nature Communications, 10(1), 5036. 10.1038/s41467-019-13003-w

Crapse, T. B., & Sommer, M. A. (2008). Corollary discharge across the animal kingdom. Nature Reviews Neuroscience, 9(8), 587–600. 10.1038/nrn2457

Flinker, A., Chang, E. F., Kirsch, H. E., Barbaro, N. M., Crone, N. E., & Knight, R. T. (2010). Single-Trial Speech Suppression of Auditory Cortex Activity in Humans. The Journal of Neuroscience, 30(49), 16643–16650. 10.1523/JNEUROSCI.1809-10.2010

Ford, J. M., & Mathalon, D. H. (2004). Electrophysiological evidence of corollary discharge dysfunction in schizophrenia during talking and thinking. Journal of Psychiatric Research, 38(1), 37–46. 10.1016/S0022-3956(03)00095-5

Ford, J. M., & Mathalon, D. H. (2005). Corollary discharge dysfunction in schizophrenia: Can it explain auditory hallucinations? International Journal of Psychophysiology, 58(2), 179–189. 10.1016/j.ijpsycho.2005.01.014

Ford, J. M., Palzes, V. A., Roach, B. J., & Mathalon, D. H. (2014). Did I Do That? Abnormal Predictive Processes in Schizophrenia When Button Pressing to Deliver a Tone. Schizophrenia Bulletin, 40(4), 804–812. 10.1093/schbul/sbt072

Ford, J. M., Roach, B. J., Faustman, W. O., & Mathalon, D. H. (2007). Synch Before You Speak: Auditory Hallucinations in Schizophrenia. American Journal of Psychiatry, 164, 458–466. 10.1176/ajp.2007.164.3.458

Galbraith, G. C. (1994). Two-channel brain-stem frequency-following responses to pure tone and missing fundamental stimuli. Electroencephalography and Clinical Neurophysiology/Evoked Potentials Section, 92(4), 321–330. 10.1016/0168-5597(94)90100-7

Galbraith, G. C., Olfman, D. M., & Huffman, T. M. (2003). Selective attention affects human brain stem frequency-following response. Neuroreport, 14(5), 735–738. 10.1097/00001756-200304150-00015

Gnanateja, G. N., Rupp, K., Llanos, F., Remick, M., Pernia, M., Sadagopan, S., Teichert, T., Abel, T. J., & Chandrasekaran, B. (2021). Frequency-Following Responses to Speech Sounds Are Highly Conserved across Species and Contain Cortical Contributions. eNeuro, 8(6), ENEURO.0451-21.2021. 10.1523/ENEURO.0451-21.2021

Greenberg, S., Marsh, J. T., Brown, W. S., & Smith, J. C. (1987). Neural temporal coding of low pitch. I. Human frequency-following responses to complex tones. Hearing Research, 25(2), 91–114. 10.1016/0378-5955(87)90083-9

Hamada, Y., Miyashita, E., & Tanaka, H. (1999). Gamma-band oscillations in the “barrel cortex” precede rat’s exploratory whisking. Neuroscience, 88(3), 667–671. 10.1016/S0306-4522(98)00468-0

Heinks-Maldonado, T. H., Mathalon, D. H., Houde, J. F., Gray, M., Faustman, W. O., & Ford, J. M. (2007). Relationship of Imprecise Corollary Discharge in Schizophrenia to Auditory Hallucinations. Archives of General Psychiatry, 64(3), 286–296. 10.1001/archpsyc.64.3.286

Houde, J. F., Nagarajan, S. S., Sekihara, K., & Merzenich, M. M. (2002). Modulation of the Auditory Cortex during Speech: An MEG Study. Journal of Cognitive Neuroscience, 14(8), 1125–1138. 10.1162/089892902760807140

Jack, B. N., Chilver, M. R., Vickery, R. M., Birznieks, I., Krstanoska-Blazeska, K., Whitford, T. J., & Griffiths, O. (2021). Movement Planning Determines Sensory Suppression: An Event-related Potential Study. Journal of Cognitive Neuroscience, 33(12), 2427–2439. 10.1162/jocn_a_01747

Kim, K. X., Dale, C. L., Ranasinghe, K. G., Kothare, H., Beagle, A. J., Lerner, H., Mizuiri, D., Gorno-Tempini, M. L., Vossel, K., Nagarajan, S. S., & Houde, J. F. (2023). Impaired Speaking-Induced Suppression in Alzheimer’s Disease. eNeuro, 10(6). 10.1523/ENEURO.0056-23.2023

Knolle, F., Schröger, E., & Kotz, S. A. (2013). Prediction errors in self- and externally-generated deviants. Biological Psychology, 92(2), 410–416. 10.1016/j.biopsycho.2012.11.017

Krieghoff, V., Waszak, F., Prinz, W., & Brass, M. (2011). Neural and behavioral correlates of intentional actions. Neuropsychologia, 49(5), 767–776. 10.1016/j.neuropsychologia.2011.01.025

Krizman, J., & Kraus, N. (2019). Analyzing the FFR: A tutorial for decoding the richness of auditory function. Hearing Research, 382, 107779. 10.1016/j.heares.2019.107779

Krizman, J., Skoe, E., & Kraus, N. (2012). Sex differences in auditory subcortical function. Clinical Neurophysiology, 123(3), 590–597. 10.1016/j.clinph.2011.07.037

Kumar, S., Dheerendra, P., Erfanian, M., Benzaquén, E., Sedley, W., Gander, P. E., Lad, M., Bamiou, D. E., & Griffiths, T. D. (2021). The Motor Basis for Misophonia. Journal of Neuroscience, 41(26), 5762–5770. 10.1523/JNEUROSCI.0261-21.2021

Liljequist, D., Elfving, B., & Roaldsen, K. S. (2019). Intraclass correlation – A discussion and demonstration of basic features. PLOS ONE, 14(7), e0219854. 10.1371/journal.pone.0219854

Martikainen, M. H., Kaneko, K., & Hari, R. (2005). Suppressed Responses to Self-triggered Sounds in the Human Auditory Cortex. Cerebral Cortex, 15(3), 299–302. 10.1093/cercor/bhh131

Musacchia, G., Sams, M., Nicol, T., & Kraus, N. (2006). Seeing speech affects acoustic information processing in the human brainstem. Experimental Brain Research, 168(1), 1–10. 10.1007/s00221-005-0071-5

Niziolek, C. A., Nagarajan, S. S., & Houde, J. F. (2013). What Does Motor Efference Copy Represent? Evidence from Speech Production. The Journal of Neuroscience, 33(41), 16110–16116. 10.1523/JNEUROSCI.2137-13.2013

Ozker, M., Yu, L., Dugan, P., Doyle, W., Friedman, D., Devinsky, O., & Flinker, A. (2024). Speech-induced suppression and vocal feedback sensitivity in human cortex. eLife, 13, RP94198. 10.7554/eLife.94198

Popovych, S., Rosjat, N., Toth, T. I., Wang, B. A., Liu, L., Abdollahi, R. O., Viswanathan, S., Grefkes, C., Fink, G. R., & Daun, S. (2016). Movement-related phase locking in the delta–theta frequency band. NeuroImage, 139, 439–449. 10.1016/j.neuroimage.2016.06.052

Price, C. N., & Bidelman, G. M. (2021). Attention reinforces human corticofugal system to aid speech perception in noise. NeuroImage, 235, 118014. 10.1016/j.neuroimage.2021.118014

Railo, H., Nokelainen, N., Savolainen, S., & Kaasinen, V. (2020). Deficits in monitoring self-produced speech in Parkinson’s disease. Clinical Neurophysiology, 131(9), 2140–2147. 10.1016/j.clinph.2020.05.038

Singla, S., Dempsey, C., Warren, R., Enikolopov, A. G., & Sawtell, N. B. (2017). A cerebellum-like circuit in the auditory system cancels responses to self-generated sounds. Nature Neuroscience, 20(7), 943–950. 10.1038/nn.4567

Sitek, K. R., Mathalon, D. H., Roach, B. J., Houde, J. F., Niziolek, C. A., & Ford, J. M. (2013). Auditory Cortex Processes Variation in Our Own Speech. PLoS ONE, 8(12), e82925. 10.1371/journal.pone.0082925

Skoe, E., & Kraus, N. (2010). Auditory Brain Stem Response to Complex Sounds: A Tutorial. Ear and Hearing, 31(3), 302. 10.1097/AUD.0b013e3181cdb272

Smith, J. C., Marsh, J. T., & Brown, W. S. (1975). Far-field recorded frequency-following responses: Evidence for the locus of brainstem sources. Electroencephalography and Clinical Neurophysiology, 39(5), 465–472. 10.1016/0013-4694(75)90047-4

Stoll, T. J., Vandjelovic, N. D., Polonenko, M. J., Li, N. R. S., Lee, A. K. C., & Maddox, R. K. (2025). The auditory brainstem response to natural speech is not affected by selective attention. PLOS Biology, 23(10), e3003407. 10.1371/journal.pbio.3003407

Suga, N., & Schlegel, P. (1972). Neural Attenuation of Responses to Emitted Sounds in Echolocating Bats. Science, 177(4043), 82–84. 10.1126/science.177.4043.82

Suga, N., Xiao, Z., Ma, X., & Ji, W. (2002). Plasticity and Corticofugal Modulation for Hearing in Adult Animals. Neuron, 36(1), 9–18. 10.1016/S0896-6273(02)00933-9

Thaut, M. H., McIntosh, G. C., & Hoemberg, V. (2015). Neurobiological foundations of neurologic music therapy: Rhythmic entrainment and the motor system. Frontiers in Psychology, 5. 10.3389/fpsyg.2014.01185

Tierney, A., & Kraus, N. (2013). The Ability to Move to a Beat Is Linked to the Consistency of Neural Responses to Sound. Journal of Neuroscience, 33(38), 14981–14988. 10.1523/JNEUROSCI.0612-13.2013

Varghese, L., Bharadwaj, H. M., & Shinn-Cunningham, B. G. (2015). Evidence against attentional state modulating scalp-recorded auditory brainstem steady-state responses. Brain Research, 1626, 146–164. 10.1016/j.brainres.2015.06.038

von Holst, E., & Mittelstaedt, H. (1950). Das Reafferenzprinzip. Naturwissenschaften, 37(20), 464–476. 10.1007/BF00622503

White-Schwoch, T., Anderson, S., Krizman, J., Nicol, T., & Kraus, N. (2019). Case studies in neuroscience: Subcortical origins of the frequency-following response. Journal of Neurophysiology, 122(2), 844–848. 10.1152/jn.00112.2019

Winer, J. A. (2005). Decoding the auditory corticofugal systems. Hearing Research, 207(1), 1–9. 10.1016/j.heares.2005.06.007

Winer, J. A., Larue, D. T., Diehl, J. J., & Hefti, B. J. (1998). Auditory cortical projections to the cat inferior colliculus. Journal of Comparative Neurology, 400(2), 147–174. 10.1002/(SICI)1096-9861(19981019)400:2%3C147::AID-CNE1%3E3.0.CO;2-9

Xie, Z., Reetzke, R., & Chandrasekaran, B. (2018). Taking Attention Away from the Auditory Modality: Context-dependent Effects on Early Sensory Encoding of Speech. Neuroscience, 384, 64–75. 10.1016/j.neuroscience.2018.05.023

